# A quantitative framework for investigating the reliability of network construction

**DOI:** 10.1101/332536

**Authors:** Alyssa R. Cirtwill, Anna Eklöf, Tomas Roslin, Kate Wootton, Dominique Gravel

## Abstract

Descriptions of ecological networks typically assume that the same interspecific interactions occur each time a community is observed. This contrasts with the known stochasticity of ecological communities: community composition, species abundances, and link structure all vary in space and time. Moreover, finite sampling generates variation in the set of interactions actually observed. Here we develop the conceptual and analytical tools needed to capture uncertainty in the estimation of pairwise interactions. To define the problem, we identify the different contributions to the uncertainty of an interaction and its implications for the estimation of network properties. We then outline a framework to quantify the uncertainty around each interaction. We illustrate this framework using the most extensively sampled network to date. We found significant uncertainty in estimates for the probability of most pairwise interactions which we could, however, limit with informative priors. Through these efforts, we demonstrate the utility of our approach and the importance of acknowledging the uncertainty inherent in network studies. Most importantly, we stress that networks are best thought of as systems constructed from random variables, the stochastic nature of which must be acknowledged for an accurate representation. Doing so will fundamentally change networks analyses and yield greater realism.

## Introduction

Representing an assemblage of species as a network offers a convenient summary of how the community is constructed as networks simultaneously describe species composition and interactions between species. A tabulation of the nodes (species) and their relative abundances forms the basis for traditional metrics of community composition such as alpha diversity. To move from these simpler metrics to a network framework, the tabulation of nodes is combined with interactions (links between nodes) so that networks provide additional, higher-order information on community structure. While this additional information is useful (as, for example, interactions can affect changes in species abundances over time), empirical descriptions of ecological networks are still limited because they are usually considered to be static representations of the communities and interactions they describe. That is, whether the network is assembled based on aggregated data, a single intensive “snapshot” sample, or expert knowledge, interactions are assumed to occur deterministically wherever and whenever the community is observed (Olesen et al., 2011).

The assumption of static communities contrasts significantly with the widely recognised stochasticity of ecological communities (Gotelli, 2000). Community composition and species abundances vary from site to site (Baiser et al., 2012) and over time within a site (Olesen et al., 2011). Likewise, interactions vary over space (Kitching & Kitching, 1987; Baiser et al., 2012), time (Kitching & Kitching, 1987; Olesen et al., 2011), and between individuals of a given species (Pires et al., 2011a; Fodrie et al., 2015; Novak & Tinker, 2015). We emphasise that variability in community composition and interactions may or may not be closely related. The removal of a species from a site will obviously also remove its interactions but, conversely, the co-occurrence of potentially interacting species does not in itself guarantee that they will interact at a given place and time. Interactions can be lost if the interaction partners remain present but are separated in time or are too rare to detect each other (Tylianakis et al., 2010). Interactions can also fail to occur because of environmental contingencies (Poisot et al., 2015), or through changes to individual preferences (Fodrie et al., 2015).

Beyond “true” variation in network structure, several researchers have pointed to the importance of sampling intensity for the assessment of network structure (e.g., Martinez et al., 1999; Blüthgen et al., 2006, 2007). An assessment of the accumulation of interactions with increasing sampling effort suggests that it is even more challenging to document interactions than species (Poisot et al., 2012). As a result, it has been proposed that interactions should be described probabilistically and network metrics computed accordingly (Poisot et al., 2016). Early work in this vein includes food-web models using likelihood-based approaches (Allesina et al., 2008) or Gaussian (Williams et al., 2010) or binomial (Rohr et al., 2016) probability functions for each possible interaction. These models may include information about species’ traits (Rohr et al., 2016) or may attempt to reproduce empirical network structures using a set of simple rules (Allesina et al., 2008; Williams et al., 2010).

Despite these preliminary efforts, to date we lack the quantitative methodology to deal with the uncertainty generated by spatiotemporal variation in ecological interactions and by sampling. Even in extremely well-sampled networks, uneven sampling across species (or pairs of species) can lead to the erroneous inference that some species do not interact because they co-occur rarely or have not yet been observed together - even if they do interact when they do co-occur (see Box 1 for an example). Nearly all network studies will thus neglect some interactions, necessitating an approach that acknowledges this uncertainty.

### Box 1

*Salix*-galler-natural enemy dataset.

As a case study, we use an extensively sampled *Salix*-galler-natural enemy meta-network. This dataset consists of a single community type sampled across Europe: willow (*Salix*) species, willow-galling sawflies, and their natural enemies. The data were collected over 29 years at 374 unique locations across Europe with a total of 641 site visits. Each site visit or each unique site can be considered as a network in its own right or as an independent sample from which to build the meta-network. Here we take the more conservative approach and pool visits to the same site for a sample size of 374 sub-networks. The meta-network consists of 1,173 different interactions between 52 *Salix* nodes, 92 herbivore nodes, and 126 natural enemy nodes. The high spatiotemporal resolution of this network and the unusually high sampling effort implemented at the site level makes this dataset particularly well suited for illustrating the difficulties in completely sampling a network and testing Bayesian approaches to overcome these difficulties.

We may begin by comparing the frequency of co-occurrences for pairs of species in each part of the network to reveal the challenge of having sufficient sampling to be confident that an interaction does not occur. Most pairs of species (3,986/4,992 *Salix*-galler pairs and 9,794/12,096 galler-natural enemy pairs) are never found co-occurring and, for species that did occur together, the total number of co-occurrences was generally low (mean=4.24, variance=36.3 for *Salix*-galler pairs; mean=3.87, variance=28.8 for galler-natural enemy pairs; Fig. 1A-B). The bulk of these co-occurring species pairs were never observed to interact: only 2.82% of *Salix*-galler pairs and 7.76% of galler-natural enemy pairs were observed interacting at one or more sites. Of those pairs that did interact, the incidence of interaction was also low (mean=12.0, variance=155 for *Salix*-galler pairs; mean=4.04, variance=29.3 for galler-natural enemy pairs; Fig. 1C-D). Thus, even in the most extensive data set that we could find, there was very little empirical data for each species pair. This suggests that limited sampling is a major source of uncertainty in all empirical networks. This dataset also illustrates the potential for increased sampling to not necessarily reveal more interactions as a pair of species that is able to interact may not be observed interacting in all samples where the pair co-occurs (Fig. 1E-F).

In this study, we formalise the description of interactions between species as probabilities and develop analytical tools to capture the uncertainty in the estimation of these interactions. We focus on binary interactions as a first step, but the framework could be expanded to deal with interaction frequencies and strength. To define the problem, we first identify the different contributions to the uncertainty of an interaction and discuss the implications of each source of uncertainty for the properties of ecological networks. Next, we develop an analytical framework to quantify the uncertainty around interactions in an empirical web. We illustrate this framework using the most extensively sampled network to date (Box 1). Finally, we offer tangible recommendations for improved descriptors of ecological interactions. Through these efforts, we demonstrate both the utility of our approach and the importance of acknowledging the uncertainty inherent in network studies.

## Why do some interactions *not* occur?

To define the problems associated with quantifying ecological interaction networks, we will start from the perspective of an empirical community ecologist faced with the task of describing a previously unknown interaction network. This ecologist will be interested in generating a description of the species/nodes present and the links between them (Roslin & Majaneva, 2016). Importantly, the information sought is conveyed by both the presence and *absence* of links. Presences and absences are not, however, equally certain. An observed link will always remain an observed link but there are multiple reasons why a given link may not be observed. Thus, the detection of any interaction is a stochastic process. We define three nested levels of uncertainty contributing to this stochasticity: interaction uncertainty, process uncertainty, and detection uncertainty.

### Interaction uncertainty

First, and most fundamentally, we do not know whether or not a pair of species have the appropriate characteristics (or traits) to interact. We define the probability of an interaction *L* given those characteristics **T** as *P*(*L*|**T**) = λ. Obviously, if *k* (the number of observed interactions) is 0, it is possible that the two species would not interact even if there were no external constraints (e.g., temporal or environmental separation) preventing the interaction from co-occurring. As a simple example, a prey species may be too large to be consumed by a particular predator. In such cases, λ would take a value of 0 and there would be no uncertainty.

Nevertheless, it is also possible that the interaction is a rare phenomenon with λ > 0 that has not yet been documented. This source of uncertainty is the one documented by trait-matching models (Bartomeus et al., 2016). It arises because every model is imperfect and lacks information (i.e. about traits) that could be used to define constraints on the interaction (Dormann et al., 2017). Further study may, however, eventually reveal the traits of interest and allow us to reduce interaction uncertainty. In other words, with sufficient sampling and all information accessible, this interaction probability λ should either tend to 0 or to 1.

### Process uncertainty

Even when an interaction is feasible, i.e. *L* = 1, it may not occur at a given location or moment in time because of local constraints such as inclement weather or the lack of suitable habitat. We define the realisation of the interaction process with the variable *X*, given that the interaction is feasible, as a stochastic process with associated probability *P*(*X* | *L* = 1) = χ. This phenomenon of interaction contingencies is usually not considered in network studies, but there is a rich literature in community ecology about the contingencies of interactions. Phenological matching (Miller-Rushing et al., 2010; Gezon et al., 2016), species preferences (Pires et al., 2011b; Novak & Tinker, 2015; Coux et al., 2016), and fear effects of other species (Luttbeg & Kerby, 2005; Wirsing & Heithaus, 2008) are just some of the factors contributing to variation in the frequency of interactions between a given pair of species. Although some of the factors leading to process uncertainty can be addressed in mesocosm studies of networks (e.g., environmental conditions can be held stable), process uncertainty is likely inevitable in the field.

### Detection uncertainty

Lastly, measurement errors are a pervasive source of uncertainty in the observation of ecological processes. Given that an interaction is feasible and occurs under the local conditions (*L*=1 and *X*=1), we may define the detection of an interaction, *D*, as a stochastic process with the associated probability *P*(*D* | *X* = 1, *L* = 1) = *δ*. Detection failure could happen for several reasons including failure to rear a parasitoid, species mis-identification, or because the interaction is very rare (see Wirta et al. (2014) for examples of some of these difficulties and partial solutions to them). Some sources of detection error can be minimised with appropriate sampling effort (δ will converge to one with increasing number of samples), but other sources are often difficult to reduce (e.g. the occurrence of cryptic species might require molecular analysis for appropriate taxonomic identification as in Wirta et al. 2014; Frost et al. 2016).

## Estimating detection and process uncertainty

Together, the combination of these three sources of uncertainty – interaction uncertainty, process uncertainty, and detection uncertainty– results in a range of potential explanations for the observation of an absence of interaction (*D*, *X*, and/or *L* = 0). The ecologist wanting to describe the network, however, is specifically interested in the situation where *L* = 0 (i.e., in true absences). Thus, while there is no difficulty interpreting the observation of an interaction, the observation of an absence of an interaction offers more of a challenge since it must be decomposed into different quantities. It is particularly important to rule out the situations where *D* = 0 ⋃ *X* = 1 ⋃ *L* = 1, i.e. where the interaction occurred at the location but was not observed, and *D* = 1 ⋃ *X* = 0 ⋃ *L* = 1, i.e., where the interaction is feasible and would have been detected but did not occur at the local site. The occurrence of a true absence, our quantity of interest, corresponds to the joint event *L* = 0 ⋃ *X* = 1 ⋃ *D* = 1 but in reality an empirical ecologist will measure the marginal probability *P*(*L*) = *k*/*n* where *k* is again the number of observed interactions and *n* the number of observed co-occurrences.

The considerations above raise a major challenge: when faced with empirical data, how may we infer whether unobserved interactions went undetected due to sampling or whether they truly do not occur? How then may we refine our sampling approaches to reduce uncertainties, and do we gain insights into the impact of multiple processes on field observations? Importantly, some sources of uncertainty can be minimised with appropriate sampling design and efforts while other sources are difficult or impossible to reduce since they are generated by chance variation created by the very process in which we are interested. Given this multifaceted problem of uncertainty, what can we do to separate the different types of variation and reduce those that can be reduced?

The obvious rule of thumb is to “sample more” (see Fig. 2 for a demonstration of the power of increasing sample size). Sampling more will clearly reduce uncertainty regarding the upper bound of the probability of interaction and it will also increase the probability of detecting unlikely interactions (e.g.,. interactions where *L*=1 but process uncertainty is high). Despite these benefits, we note that there are limits to the utility of increased sampling. Since the probability of observing the co-occurrence of two species will always be higher than the probability of observing their interaction (since the probability of interaction is conditional on both interaction partners being present; see Fig. 1E-F), we will accumulate observations of co-occurrences faster than we will accumulate observations of interactions. Thus, the more we sample, the more zeros will appear in our interaction matrix.

**Figure 1:**
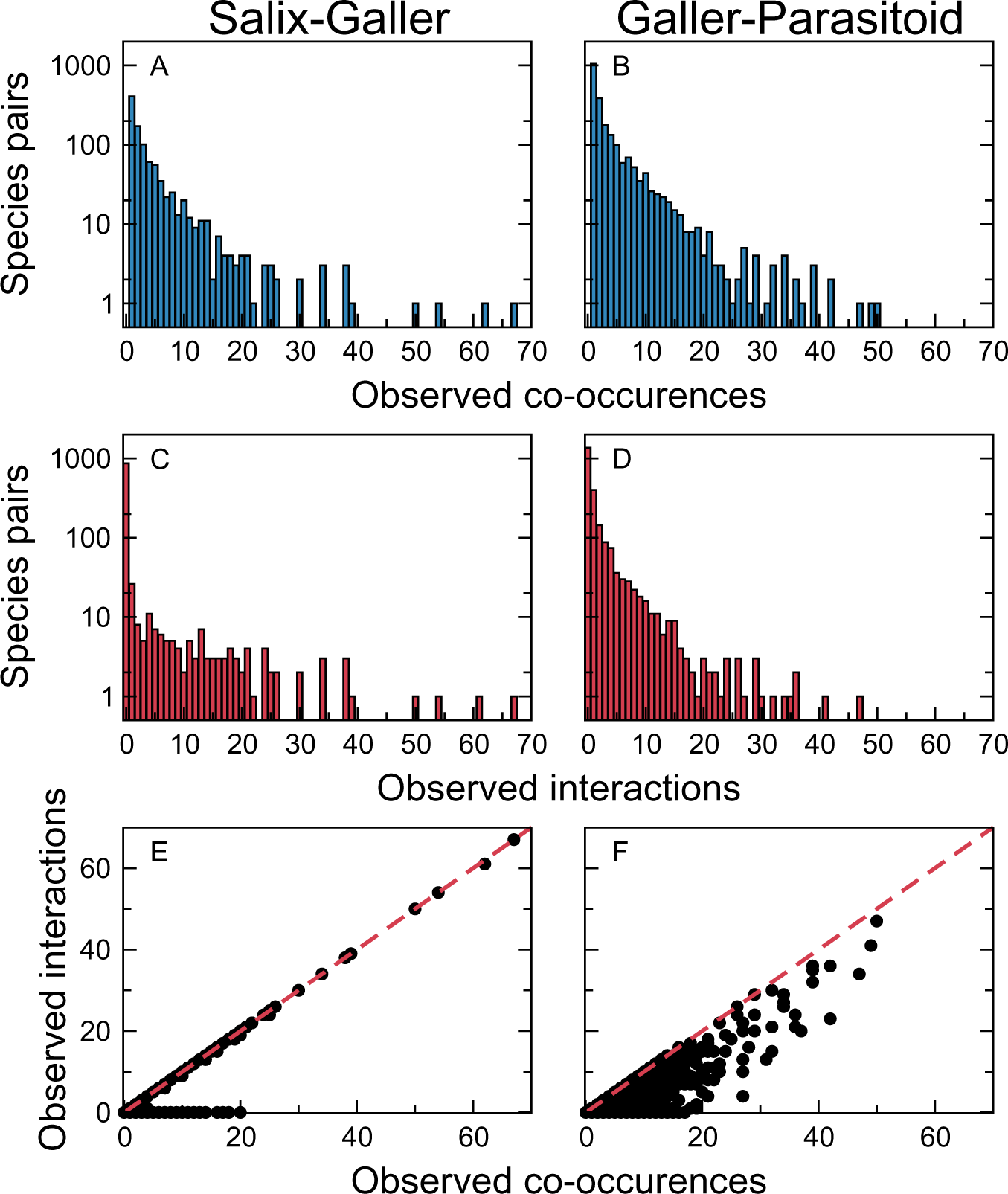
**A-B)** Most pairs of *Salix* and gallers or gallers and natural enemies were never observed co-occurring despite the high levels of replication in our example dataset. For those pairs that were observed together at least once (*n_ij_* > 0), the number of observed co-occurrences was generally small (<10). Here we show a histogram of the number of pairs of species observed co-occurring at least once. 3986 *Salix*-galler and 9794 galler-enemy pairs were never observed co-occurring: these pairs are omitted from the histogram. **C-D)** Most pairs of species that were observed at the same site were never observed interacting. Here we show a histogram of the number of observed interactions within pairs of co-occurring species. Species which co-occurred but never interacted are included in these histograms. **E-F)** Here we show, for each species pair, the number of observed interactions plotted against the number of observed co-occurrences. *Salix*-galler pairs either are never observed interacting or interact almost every time they co-occur, while galler-enemy pairs had more variable frequencies of interaction. In panels E and F the red, dashed line indicates a 1:1 relationship between interactions and co-occurrences.

**Figure 2:**
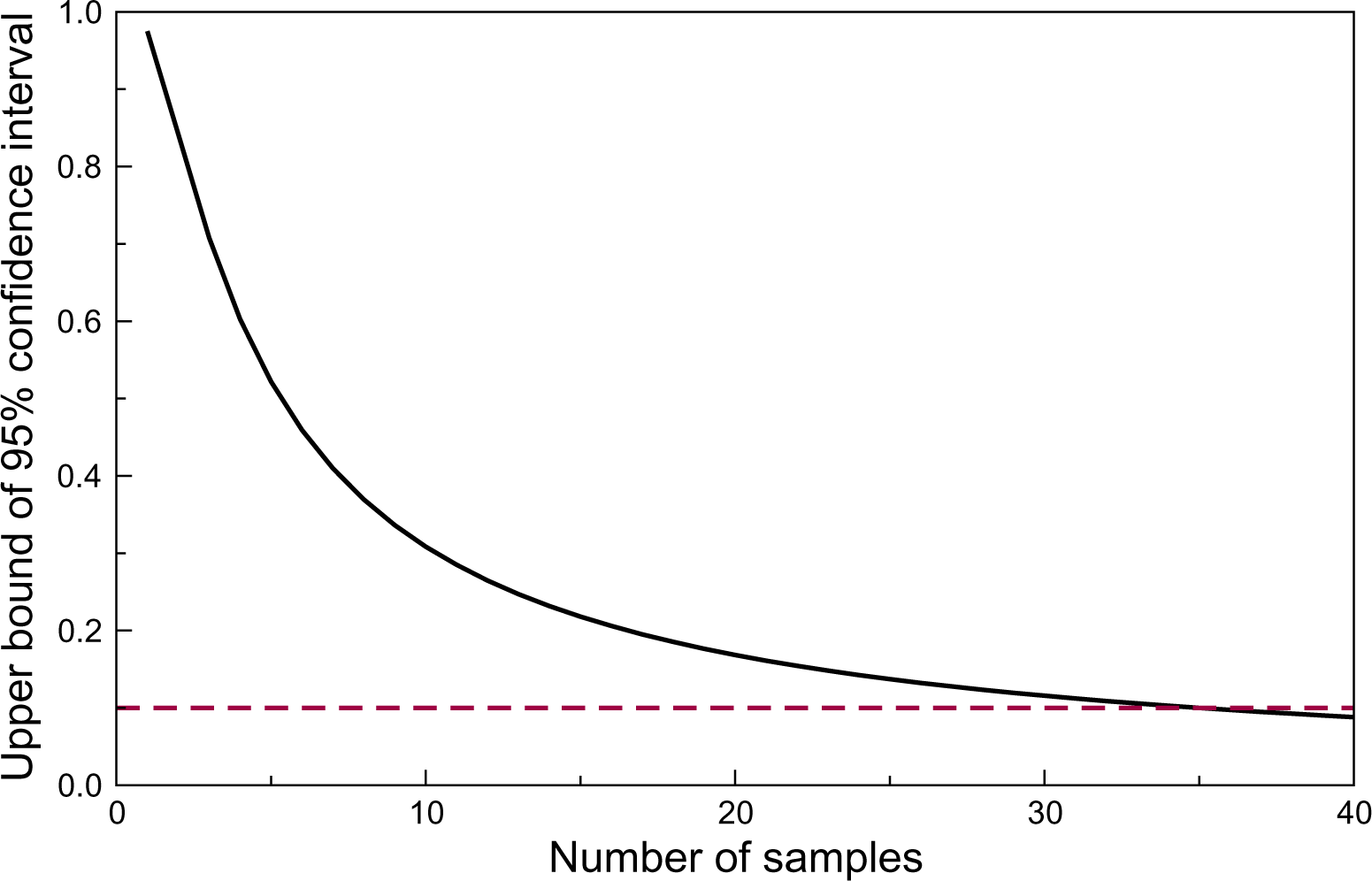
A simple example will illustrate the problem of imperfect detection of interactions. Assume that we want to infer the probability of an interaction between two species, *i* and *j*. Now assume that in reality, interaction between *i* and *j* is completely impossible (i.e. the true λ = 0) but the observer does not know this and seeks to estimate this interaction probability (λ). The number of observed interactions will follow a binomial distribution with number of interactions *k* and number of observations *n*. Using this distribution, we can compute the credible interval of the estimated probability λ. Even assuming no added detection error in observing the incidence of the interaction, a single observation of species co-occurrence reveals very little regarding the probability of the interaction as the credible interval for a pair of species with one observation essentially spans from 0 to 1. Only with 35 observations will the upper limit of the credible interval be lowered to 0.1. Thus, adding more observations is certainly useful in controlling uncertainty, but the number of observations added needs to be very high. Here we show the upper bound (solid black line) of a 95% Clopper-Pearson true credible interval for λ when *k* = 0 (*i* and *j* have not been observed interacting) for a variety of *n* (observed co-occurrences of *i* and *j*). Using a Bayesian approach with an informative prior can reduce the confidence interval about λ for a given sample size. λ threshold interaction probability of 0.1 is indicated by the dashed red line.

In one endeavour to determine whether unobserved interactions were undetected due to sampling, or whether they truly do not occur, Weinstein & Graham (2017) used repeated sampling rounds to estimate the daily probability of detecting a hummingbird interaction, and to thereby model detection and process uncertainty. While conceptually attractive, this approach is unsuitable for interactions occurring over longer time scales (e.g., associations between hosts and parasitoids with a single generation per year), or very rare interactions which might not occur on any of the sampling days or might involve individuals of a species that is not under observation. What is worse, the problem persists that the absence of an interaction of a given day could either be because it was impossible on that day despite being otherwise feasible [*P*(*X* | *D* = 1, *L* = 1) = 0], because interaction did occur but could not be observed [*P*(*D* | *X* = 1, *L* = 1) = 0], or any combination of the two. From a conceptual perspective, this approach therefore fails to satisfactorily distinguish between sources of uncertainty. Most importantly, if two species are never observed co-occurring during several days of sampling then we have learned nothing about their probability of interacting if they should ever co-occur. In other words, there is no information about interactions without co-occurrence.

An added complication is that not all sources of uncertainty are proportional to sample size. To record an interaction between A and B, we need to identify both partners correctly (a non-trivial problem in many food webs; e.g. Kaartinen & Roslin, 2011; Roslin & Majaneva, 2016) and be able to resolve all interactions with a similar likelihood. For both molecular and rearing techniques, certain types of interactions may go unnoticed due to technical challenges (Wirta et al., 2014). This can bias the set of recorded interactions. The bottom line is that separating different sources of uncertainty is difficult indeed. As an alternative to abandoning empirical networks or continuing to ignore the uncertainty inherent in undetected observations, we propose that some insight regarding the detectability of interactions between species not found co-occurring in a focal system may be gained from data on other species pairs in the same or a similar system.

## A naive quantification of uncertainty

To progressively dissect the different contributions to uncertainty, we will start by considering how we could naively quantify interaction probability and its associated uncertainty *for an interaction that has not yet been observed*. We consider the case where a pair of species have been observed co-occurring *n* times, of which they have been observed to interact in *k* = 0 cases. We now aim to evaluate the uncertainty of this interaction. We consider the occurrence of an interaction as a Bernoulli trial. Consequently, the number of successes *k* over *n* trials will follow a binomial distribution:

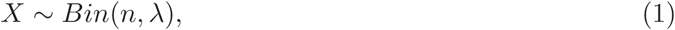

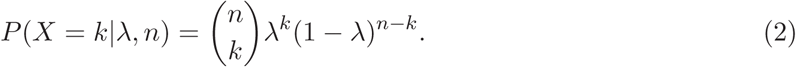

The parameter λ, the probability of observing an interaction over an infinite time interval and area, is the quantity we want to estimate from empirical data. The maximal likelihood estimate (MLE) of λ is straightforward to find given *k* and *n*:

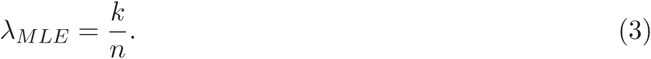

The variance of a Bernoulli experiment is *n*λ(1-λ). It is important to remember that this variance describes the variability of the number of successes *k* for *n* trials and is not the variance associated with the estimation of λ. Given this variance, it is possible to compute the confidence interval for the MLE of λ using any of several methods, including the *Wilson score interval*, the *Clopper-Pearson interval*, and the *Agresti-Coull interval* (for details, see [Brown et al., 2001]). Finding this estimate is therefore quite straightforward, but it nonetheless has two drawbacks. First, λ is not a single point estimate but rather a random variable with an unknown distribution. This means that if *k* = 0 in a given sample, this does not necessarily imply that the two species will never interact. Rather, *k* = 0 implies that ‘no interaction’ is the most likely outcome when the species do co-occur but there is nonetheless a substantial chance that the two species *could* interact. In the situation where *k* > 0, in contrast, we are sure that the interaction is feasible (*L* = 1) but still cannot be sure of the cause if the interaction is not observed at some sites/times (i.e., we cannot say why *k* < *n*). There may be local constraints (*X* = 0) or we might simply not observe the interaction in every sample (*D* < 1).

Second, where the number of samples *n* is very low (some pairs of species may never have been documented as co-occurring), there will be considerable uncertainty around our estimate of λ. In Fig. 2 and Box 2, we derive the Clopper-Pearson interval to explore how the estimate of λ varies with sample size. At a small sample size, the 95% confidence interval spans all values of λ. To establish that species are not interacting with any acceptable certainty requires tens of observations of the two species co-occurring but not interacting. As most data sets will lack such extensive sampling across all species pairs, we can use a Bayesian approach to supplement what data we do have with other sources of information.

### Box 2

Calculating the credible interval around a probability estimate

Here we describe the derivation of the Clopper-Pearson credible interval for the estimated probability of interaction λ of a pair of species observed co-occurring *n* times and interacting *k* times. As we are most interested in the probability of interaction between species pairs that have never been observed co-occurring, we consider only the case where *k* = 0 over a variety of *n*. This is straightforward to do in R (see the function “credible_interval” in *Appendix S1*).

First, we must obtain the *α* and *β* parameters for the prior distribution. In this study we obtained these parameters using the R (R Core Team, 2016) function fitdist from the package fitdistrplus (Delignette-Muller & Dutang, 2015). Once *α* and *β* are known, we can update them using our observed data. Specifically, we are interested in *α*′ = *α* + *k* and *β*′ = *β* + *n* − *k*. These parameters can then be used to calculate a credible interval using the R (R Core Team, 2016) function qbeta. In the table below, we present the 95% credible intervals for *Salix*-galler and galler-natural enemy pairs with different numbers of observed co-occurrences (*n*) and no observed interactions (*k* = 0), calculated using prior information derived from the Zillis sub-network (Kopelke et al., 2017).

**Table 1:**
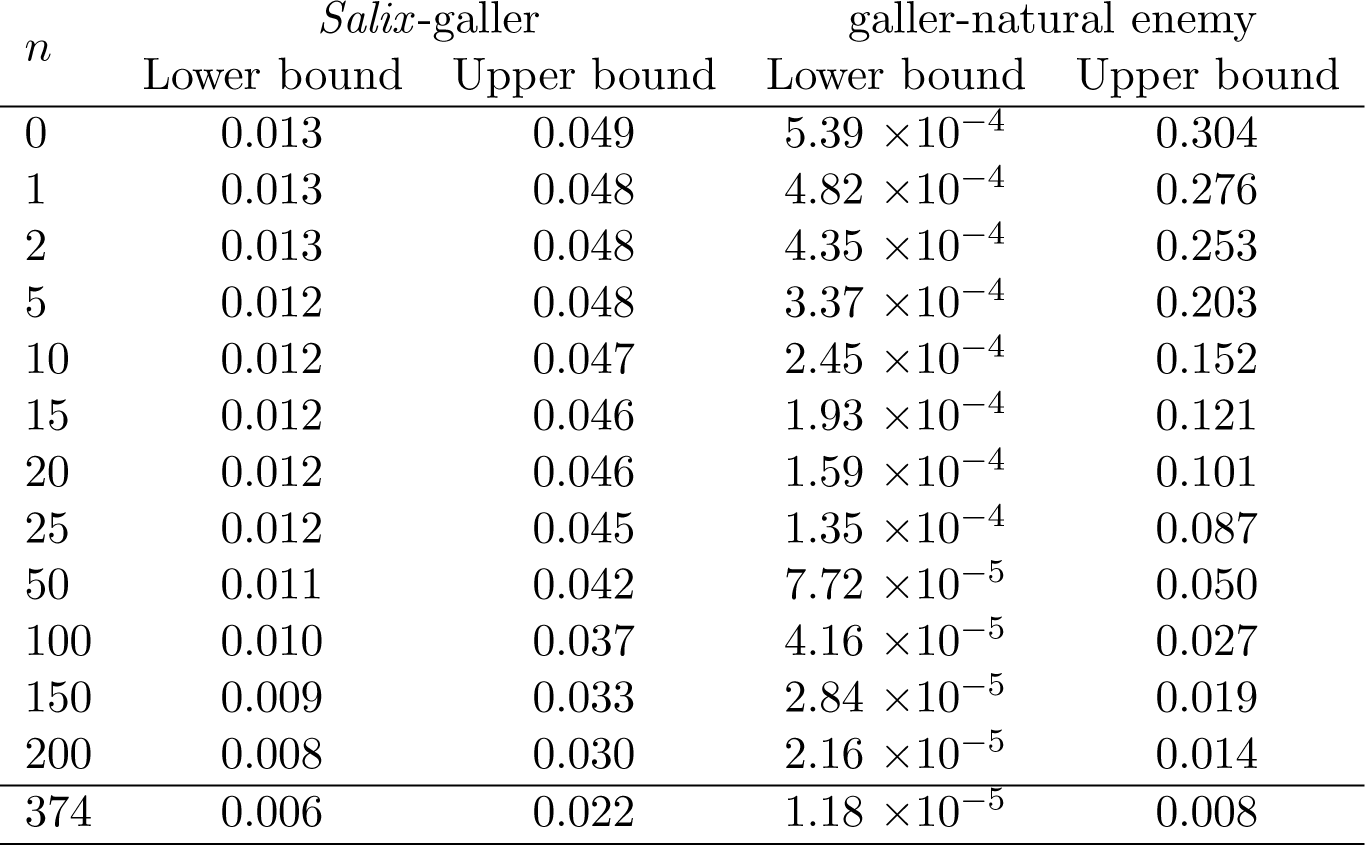
Here we give the lower and upper bounds of 95% credible intervals for the probability of interaction λ between *Salix*-galler or galler-natural enemy pairs that have been observed co-occurring *n* times but have never been observed interacting.

## Bayesian approach to infer interaction probabilities

### Posterior distribution of the interaction probability

Here we adopt a Bayesian approach to estimate the posterior distribution of the parameter λ:

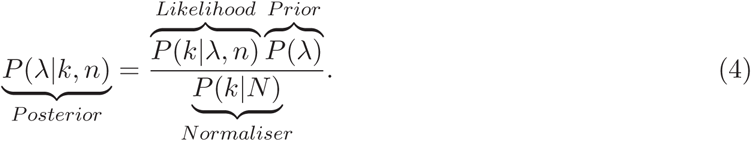

According to the above description, the likelihood is simply the binomial distribution (Eq. 2). Since λ is a probability, it is bounded between 0 and 1 and the most appropriate prior distribution is the beta:

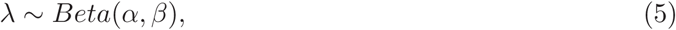

which has two shape parameters, *α* and *β*.

The beta-binomial distribution is a conjugate distribution of the binomial distribution. This allows us to analytically compute the posterior distribution of a binomial model with a beta prior distribution. We can re-write the posterior distribution of λ as:

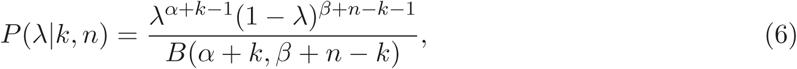

where the function *B* is the beta function. The posterior distribution of λ therefore follows the beta distribution with new parameters *α*′ = *α* + *k* and *β*′ = *β* + *n* − *k*. The weight of the prior on the posterior distribution can be understood from these parameter definitions: the difference between the posterior and the prior will increase with *k* and *n* − *k*. In other words, the distribution of λ for better-sampled pairs of species will rely less on the information used to build the prior distribution and depend more on the observed data. When plotted, we find the shape of the distribution gets narrower with *k* and *n* (Fig. 3).

**Figure 3:**
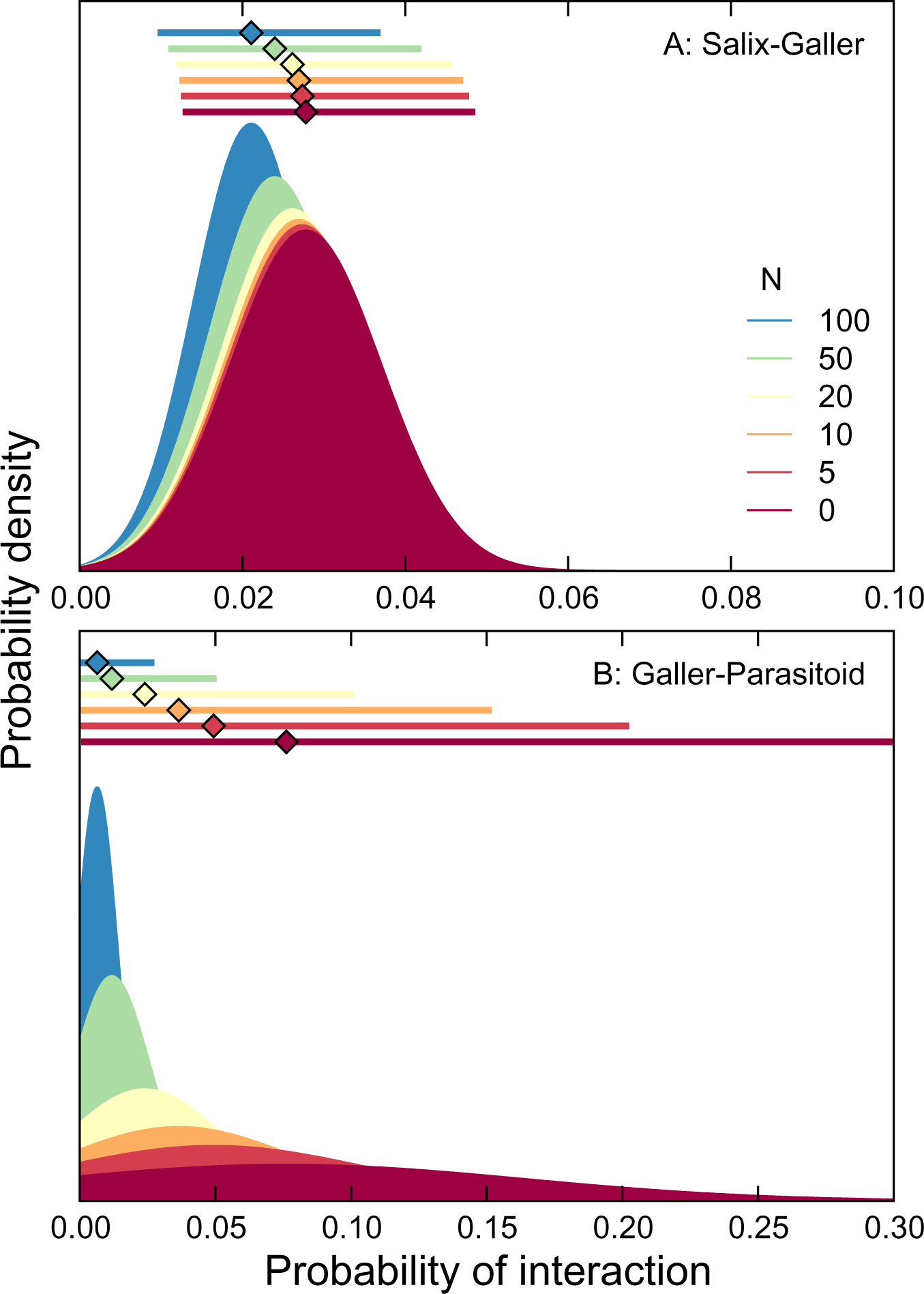
Using prior distributions based on the *Salix*-galler and galler-natural enemy networks sampled at a single site in Kopelke et al. (2017), we can calculate posterior dis-tributions for the probability of interaction (λ) between two species that have not yet been observed interacting. Here we show posterior distributions for λ in each network component ranging from the prior distribution (*n* = 0 observed co-occurrence) to the distribution obtained when the pair of species has been observed co-occurring 100 times. The distribution narrows and approaches zero as the sample size increases. Likewise, the maximum likelihood estimator for the mean probability of interaction (diamonds at top of each panel) approaches zero and the 95% credible interval (lines at top of each panel) narrows as sample size increases. **A**) The posterior distributions for λ in the *Salix*-galler component are narrower at low *n* but shrink less with increased sampling than those for **B**) the distributions of λ in the galler-natural enemy component.

### Moments and other properties

It is common to preform analyses that require calculating higher-order network properties in interaction networks. The fact that the posterior distribution of λ follows a beta distribution makes it straightforward to compute moments and other properties needed for this.

The **average** of λ is:

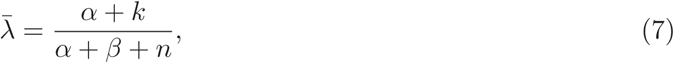

and its **variance** is:

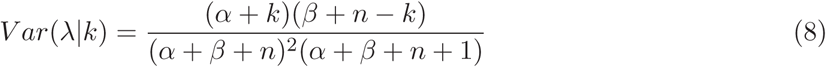

The **mode** of the distribution is:

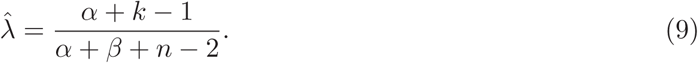

### The prior distribution

Parameters *α* and *β* determine the shape of the prior distribution, which follows a beta distribution. These are called hyper parameters. Below we identify four ways to formulate the prior distribution of λ.

### Uninformative prior

In the absence of any external information, an uninformative prior is the most conservative hypothesis for the distribution of λ. The beta distribution is in this case α uniform distribution, specified with hyper parameters *α* = 1 and *β* = 1.

### Distribution of connectance

The ecological network literature boasts a collection of networks for which connectance has been calculated and for which we can thus define the connectance distribution. Connectance is measured as *C* = *L*/*S*^2^, where *L* is the number of interactions and *S* is the number of species. It measures the filling of an interaction matrix and thereby expresses the average probability that any two species interact with each other. If we know only the mean 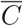 and the variance 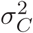 of the distribution of *C*, then the beta parameters could be computed as follows using the method of moments:

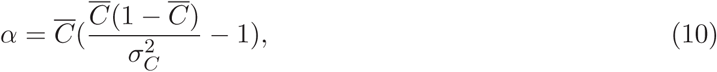

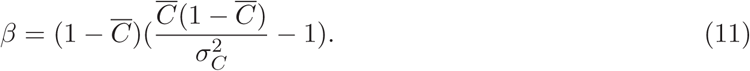

### Degree distribution or interaction probabilities

The degree of a node in a network is defined as its number of connections to other nodes. The degree distribution of a network is then the probability distribution of these degrees over the whole network and the standardised degree could therefore be interpreted as an interaction probability. It is consequently possible to use the degree distribution to inform the prior distribution. The degree distribution could come from several networks, from a similar network (e.g. a known network at slightly different location) or from the network of interest if interaction probabilities for some species are already documented. The latter approach allows researchers to apply information from known, abundant species to the rarest species for which interactions are less frequently documented.

If our focal network describes a system similar to that in a known network, we can use the distribution of interaction probabilities in that network to inform our prior. The probability of any interaction *L_ij_* depends on the degrees of species *i* and *j*. Using normalised degrees Δ_*i*_ and Δ_*j*_ (i.e., degrees divided by the number of species in the network), we can obtain the probability of interaction *L_ij_*=Δ_*i*_ × Δ_*j*_. Similar to the procedure for degree distribution, the distribution of these interaction probabilities can be used to establish a prior distribution before any data from the focal network are collected. For distributions of either degrees or interaction probabilities, the procedure for the estimation of the hyper parameters follows the same approach as described above for connectance except that each measurement is at the individual interaction level instead of the network level.

### Trait-matching function

As a fourth and final approach, it may be possible to obtain the prior distribution of λ using the outcome of a trait-matching model, provided such a model has been parameterised using external data and relevant traits are available. In such a case, the prior distribution would follow the function *P*(λ|**T**) = *f* (**T**) based on a set of traits for both species **T**. There are several techniques available to perform this inference of interaction probability, some of which are Bayesian, and we refer to Bartomeus et al. (2016) and Weinstein & Graham (2017) for recent reviews about this topic. Note that in this case the prior might not be beta-distributed and numerical methods might be required to compute the posterior distribution.

### A quantitative example

The Bayesian framework can be illustrated with a simple quantitative example. Suppose we have *n* = 10 observations of co-occurrence between species *i* and species *j* in a given time interval and area, and *k* = 3 observations of interactions. The maximum likelihood estimate of the interaction probability is simply λ_*MLE*_ = 3/10 = 0.3.

Now consider we know that species *i* is known to interact with 10 species (other than species *j*), which have the following degrees:

> *degree=c(14, 4, 2, 3, 17, 6, 2, 15, 1, 1)*.

If the network has 20 species total, this gives the normalised degrees:

> *norm_degree=c(0.65, 0.20, 0.10, 0.15, 0.85, 0.30, 0.10, 0.75, 0.05, 0.05)*.

Species *i* has a normalised degree of 0.55 (it interacts with species *j* and 10 other species). We can combine the normalised degree of *i* with the normalised degrees of its interaction partners to obtain the following set of interaction probabilities for species *i* and each of its interaction partners:

> *int_probs = c(0.358, 0.110, 0.055, 0.082, 0.468, 0.165, 0.055, 0.412, 0.028, 0.028)*.

The mean of these interaction probabilities is 0.176, approximately two-thirds the λ_*MLE*_ obtained from the observed data. We can use the distribution of these interaction probabilities as our prior distribution and estimate the uncertainty surrounding our λ_*MLE*_. With some simple R code (function “calculate_parameters”, *Appendix S1*), we obtain prior parameters *α*=0.998 and *β*=4.63. Using these priors in equations 7 and 8 above (or in the R function “calculate_distribution” in *Appendix S1*), we find a prior 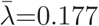 and var(λ)=0.026. Adding the observed data (*n* = 10, *k* = 3) and using the same code, we obtain posterior parameters *α*′=4.00 and *β*′=11.6 and a posterior 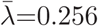=0.256 and var(λ)=0.012. Comparing the posterior distribution to the prior, we see that the posterior is closer to the observed data and that the additional data about interactions between species *i* and *j* has reduced the variance. We may also wish to calculate a credible interval (analogous to the frequentist confidence interval). This is also quite straightforward in R (see function “credible_interval” in *Appendix S1*). In this case, a 95% credible interval for 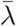 is (0.080, 0.491).

Now, consider the case where the two species have never been observed interacting across *n* trials, i-e. *k* = 0. The question is then “what is the probability that these two species do not interact”? Since it is not possible to prove that the two species could never interact (strictly speaking, in a Bayesian approach λ = 0 is impossible), we must fix a threshold below which we consider that there is no interaction (λ ~ 0). We call this threshold probability λ*. We then use the cumulative distribution function to estimate *P*(λ < λ * |L = 0, *n*) for different *n*. The function “samples_for_threshold” in *Appendix S1* calculates distribution function for λ* with an increasing number of trials. This yields a surprising result: it requires >24 observations of no interactions to be 95% sure that the interaction probability is smaller than λ*=0.1 (recall Fig. 2, Box 2). Note the special case where there is no observation of the two species co-occurring, *n* = 0. In this situation, the posterior distribution converges to the prior distribution since the data include no information on the probability with which species might interact should they co-occur.

## Scaling up to networks - an empirical example

In the following section, we will provide an empirical example based on the well-sampled system of *Salix* plants, herbivorous gallers, and their natural enemies described by Kopelke et al. (2017); see Box 1 or *Appendix S2* for a description). Using this dataset, we will demonstrate the derivation of prior distributions for the *Salix*-galler and galler-natural enemy components of these networks and the differences between these priors and posterior distributions which include all information available in this dataset (Kopelke et al., 2017). Finally, we will calculate network properties using a suite of networks sampled from these posterior distributions and show how the uncertainty around interactions that have not been observed impact these metrics.

### Computing the posterior distribution

In a strict Bayesian framework, we wish to use a prior distribution that does not rely on any information from the study at hand. Network data for a similar study system may, however, not be available. In that case, one might use the first sub-network collected as “training data” to guide future sampling. To simulate this situation, we created priors using a single sub-network from the middle of the geographical distribution of the Kopelke et al. (2017) dataset. To demonstrate how the use of data from a different system can affect the prior distribution and conclusions based on it, we repeated our analyses using priors derived from a much smaller *Salix*-galler-natural enemy system (Barbour et al., 2016, Data available from the Dryad Digital Repository: https://doi.org/10.5061/dryad.g7805). This smaller system was much more densely-connected than that described in Kopelke et al. (2017) and provided unreasonable distributions for interaction probabilities (*Appendix S4*).

To obtain the priors based on the Zillis sub-network, we estimated frequencies of *Salix*-galler interactions based on the normalised degree of each species in each network component (see *Appendix S3* for details and code). Specifically, we obtained prior parameters of *α*=8.72, *β*=305 for the *Salix*-galler component and *α*=0.700, *β*=8.49 for the galler-natural enemy components of the network. After calculating these prior parameters, we were then able to estimate the posterior distribution of interaction probabilities given the additional information in our dataset.

For species where no co-occurrences were observed (*n* = 0), we can calculate the estimates for the mean and variance of λ_*ij*_ directly from the prior parameters following equations 7 and 8 (see *Appendix S1* for R implementation). For the *Salix*-galler network, the prior distribution was: 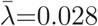=0.028, var(λ)=8.60×10^−5^. The prior distribution for the galler-natural enemy network was: 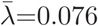=0.076, var(λ)=0.008. The posterior interaction probabilities obtained based on the Zillis sub-network were much lower than those obtained based on Barbour et al. (2016, Data available from the Dryad Digital Repository: https://doi.org/10.5061/dryad.g7805); this emphasises the importance of using an appropriate study system when constructing a prior (*Appendix S4*).

For a pair of species with some observed co-occurrences (*n* > 0), we can update the prior distribution with these data. If we consider only pairs of species which were observed to co-occur but not to interact, *k_ij_* is always 0 and only *n_ij_* will vary between species pairs, giving *α*′ = *α* and *β*′ = *β* + *n_ij_*. As the most extreme case, consider a pair of species which co-occurred at all 374 sites and was never observed to interact. Using the priors described above, our distribution for the *Salix*-galler network would become 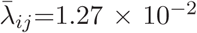, var(λ_*ij*_)=1.82 × 10^−5^ while our distribution for the galler-natural enemy network would become 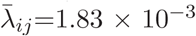, var(λ_ij_)=4.76. Distributions for both network components were very close to 0 with small variance about our estimate of λ; species *i* and *j* are extremely unlikely to interact at sites or times not included in our sample.

For most pairs of species *i* and *j*, however, *n_ij_* was much less than 374 and our posterior mean and variance therefore retain more of the influence of the prior. We can see this in the increasing means and variances as we decrease *n_ij_* (Fig. 3). The change in distribution as *n_ij_* decreases can also be shown by calculating 95% credible intervals for λ (see the function “credible_interval” in *Appendix S2*). The 95% credible interval around the estimate of λ also widens as *n_ij_* decreases from (0.001, 0.017) and (<0.001, 0.11) for hypothetical *Salix*-galler and galler-natural enemy pairs that might be observed co-occurring at all 374 sites without any observed interaction to (0.152, 0.931) and (0.008, 0.364) for *Salix*-galler and galler-natural enemy pairs that were never observed co-occurring. The 95% credible interval for hypothetical *Salix*-galler pairs widened from (0.006, 0.022) if the pair co-occurred at all sites to (0.013, 0.049) if they co-occurred at none. The 95% credible interval for hypothetical galler-natural enemy pairs, meanwhile, widened from (0.00001, 0.008) to (0.0005, 0.304).

### How many samples are required to reach a minimal precision

Rather than calculating credible intervals for a posterior distribution after collecting data, we may wish to know how many data points are necessary to obtain a given level of confidence that two co-occurring species do not interact. The number of samples needed will depend on both our desired level of confidence and the threshold below which we assume that two species are unlikely to ever interact (Fig. 4; see function samples_for_threshold in *Appendix S1*). In our dataset, the entire 95% credible interval was (0.013, 0.049). We may therefore be 95% confident that the interaction probability for *Salix* and galler species that have not been observed co-occurring is below 0.05. As the peak of the prior distribution for the probability of interaction between *Salix* and galler probabilities is around 0.02 (Fig. 3), to be 95% confident that the interaction probability for these species is below 0.01 would require 1029 observed co-occurrences with no interaction - far more than the number of sites in the (Kopelke et al., 2017) dataset.

**Figure 4:**
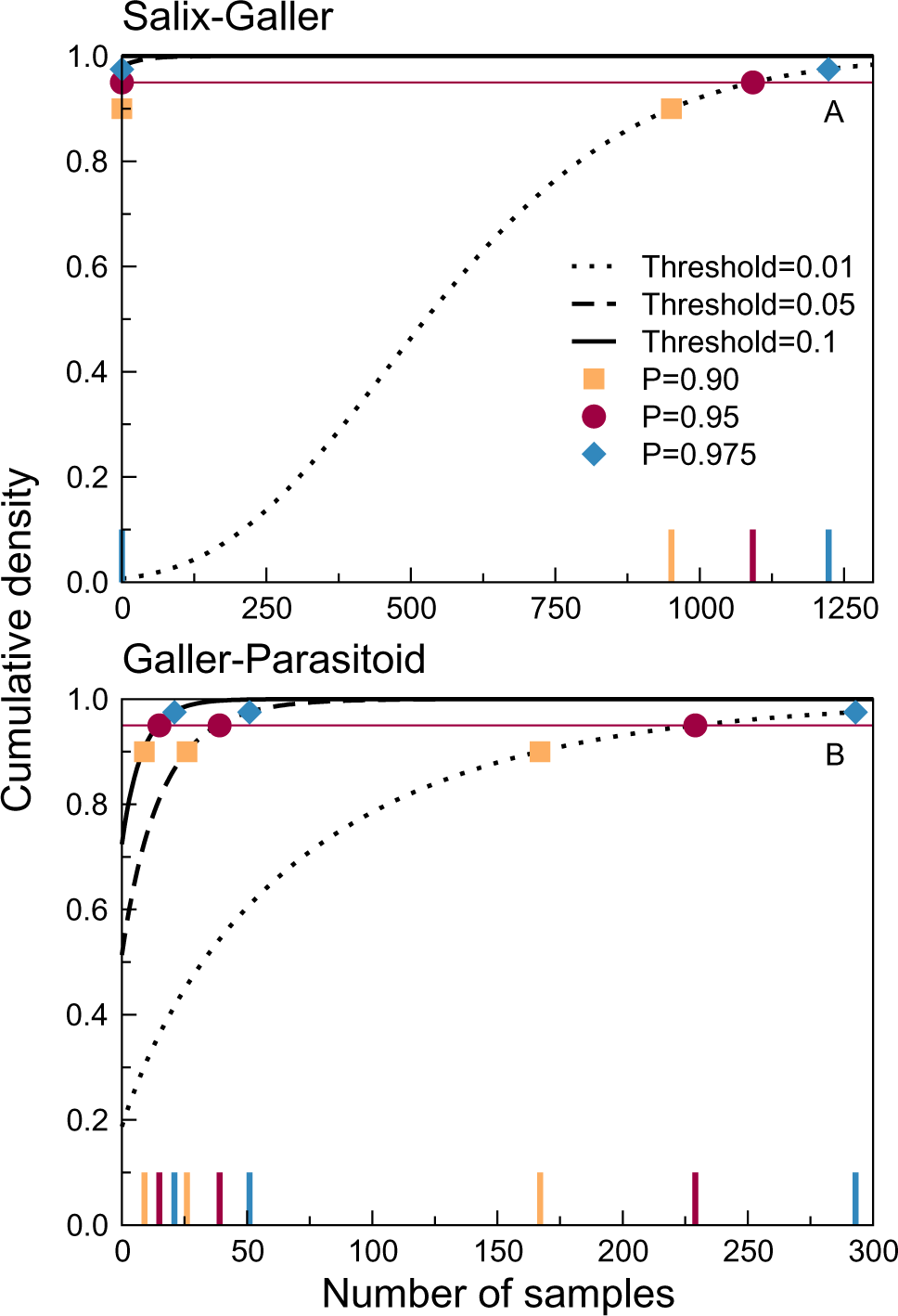
The number of samples required to achieve a given level of confidence that an interaction probability λ_*ij*_ is below a given threshold varies with both parameters. With a low threshold, our confidence that λ_*ij*_ is below the threshold increases rapidly with repeated observation of co-occurrence without interaction. Here we show the cumulative density functions for threshold probabilities of 0.5 (solid line), 0.25 (dashed line), 0.1 (dash-dot line), and 0.05 (dotted line) as well as the points at which the cdf reaches 0.90 (orange square), 0.95 (red circle), and 0.975 (blue diamond) for each threshold value. The large ticks along the x-axis indicate the number of samples associated with each of these points. **A)** In the *Salix*-galler network component, the 95% credible interval for λ_*ij*_ when *n*=0 was (0.013, 0.049). We can therefore be at least 95% confident that λ_*ij*_ is below thresholds of 0.1 or 0.05 without any observed co-occurrence of species *i* and *j*. To be confident that λ_*ij*_ is less than 0.01, however, would require more observed co-occurrences than there are sites in our dataset. **B)** In the galler-parasitoid network component, the 95% credible interval for λ_*ij*_ was substantially broader and many observed co-occurrences (≈ 15-35) are required to be 95% confident that λ_*ij*_ is below thresholds of 0.1 or 0.05.

The number of samples required to be 95% confident that the interaction probability between galler and natural enemy species is below a threshold also increases quickly as the threshold decreases. The 95% credible interval is (<0.001, 0.303) for the probability of interaction between two species observed to co-occur but never interact. To be 95% confident that the probability of interaction is below 0.1, 0.05, or 0.01 would require 15, 39, and 229 observed co-occurrences, respectively.

Given the low levels of replication in most network studies, this implies that we should have fairly low confidence in many “non-interacting” pairs of species. Even in the extensively replicated *Salix*-galler-natural enemy dataset, very few species pairs were observed co-occurring frequently enough to reach these thresholds. Regardless of our choice of prior, no species pairs were observed to co-occur frequently enough to reach the threshold for an interaction probability of 0.01. Discounting potential interactions, then, requires either a stronger prior expectation of no interaction (e.g. for forbidden interactions) or very extensive sampling. For all we know, most links absent from current descriptions of network structure may be so not because the species do not interact, but because we have not sampled deeply enough to detect them.

### Scaling up to network metrics

It is fairly straightforward to compute most network metrics when the different λ of the adjacency matrix are known and assumed not to vary without variance (Poisot et al., 2016). Several of these metrics derive directly from quantitative indices of network structure which are equivalent to λ. The remainder, originally defined for binary networks, can be adjusted to account for interaction probabilities between zero and one. It is not as easy, however, to understand how the uncertainty in these estimated interaction probabilities influences network metrics. Computation of these metrics involves non-linear functions. Since Jensen’s inequality states that the average of a non-linear function of a stochastic variable differs from the function of the average of that variable, any uncertainty in the values of λ could bias both the mean and variance of a network metric. One way to avoid potentially biased analytical calculation of network properties is to calculate the properties of a suite of simulated networks.

Using the prior distributions and procedures described above, we calculated posterior probability distributions for *Salix*-galler or galler-natural enemy pairs that were not observed interacting. Using these posterior distributions and assuming probabilities of 1 for pairs of species that were observed interacting, we created a suite of 100 webs of each network type by randomly sampling from each posterior distribution. After obtaining these posterior networks, we calculated the connectance of each web, as well as the number of links per resource (*Salix* in the *Salix*-galler networks or galler in the galler-natural enemy networks) and links per consumer. To demonstrate how these network metrics will be affected by detection uncertainty, we then created a suite of filtered networks for each posterior network. Networks were filtered by randomly sampling 99%, 95%, 90%, 80%, 70%, 60%, and 50% of the interactions included in each posterior network. This gradient is akin to a gradient of sampling effort. For each level of detection accuracy, we created 100 randomly-sampled networks per posterior-probability network (giving 100 posterior networks and 1000 detection-filtered networks each for the *Salix*-galler and galler-natural enemy networks). We then calculated the same network properties as described above.

We find, perhaps not surprisingly, that the posterior webs for the Salix-galler network had higher connectances than the original, observed web (C=0.028 for the observed web and 0.082 ≤ C ≤ 0.096 for the posterior webs; Fig. 5A). The number of links per *Salix* species in the observed web (*L_salix_*=2.71) was similar to those in the posterior webs (2.53 ≤ *L_salix_* ≤ 3.19; Fig 5C). The number of links per galler, however, was lower in the observed web (*L_galler_*=1.47) than in the posterior webs, accounting for the increased connectance (4.67 ≤ *L_galler_* ≤ 5.88; Fig. 5E). There was a more substantial difference in the nestedness of the observed and posterior webs: the observed network had NODF=0.560 while the posterior networks were more nested (1.39 ≤ NODF ≤ 1.94). Even the networks sampled with a detection filter of 50% had non-zero nestedness (Fig. 5G). This last result highlights the potential for the possibility for network structure to vary when considering the possibility that unobserved species pairs may interact.

**Figure 5:**
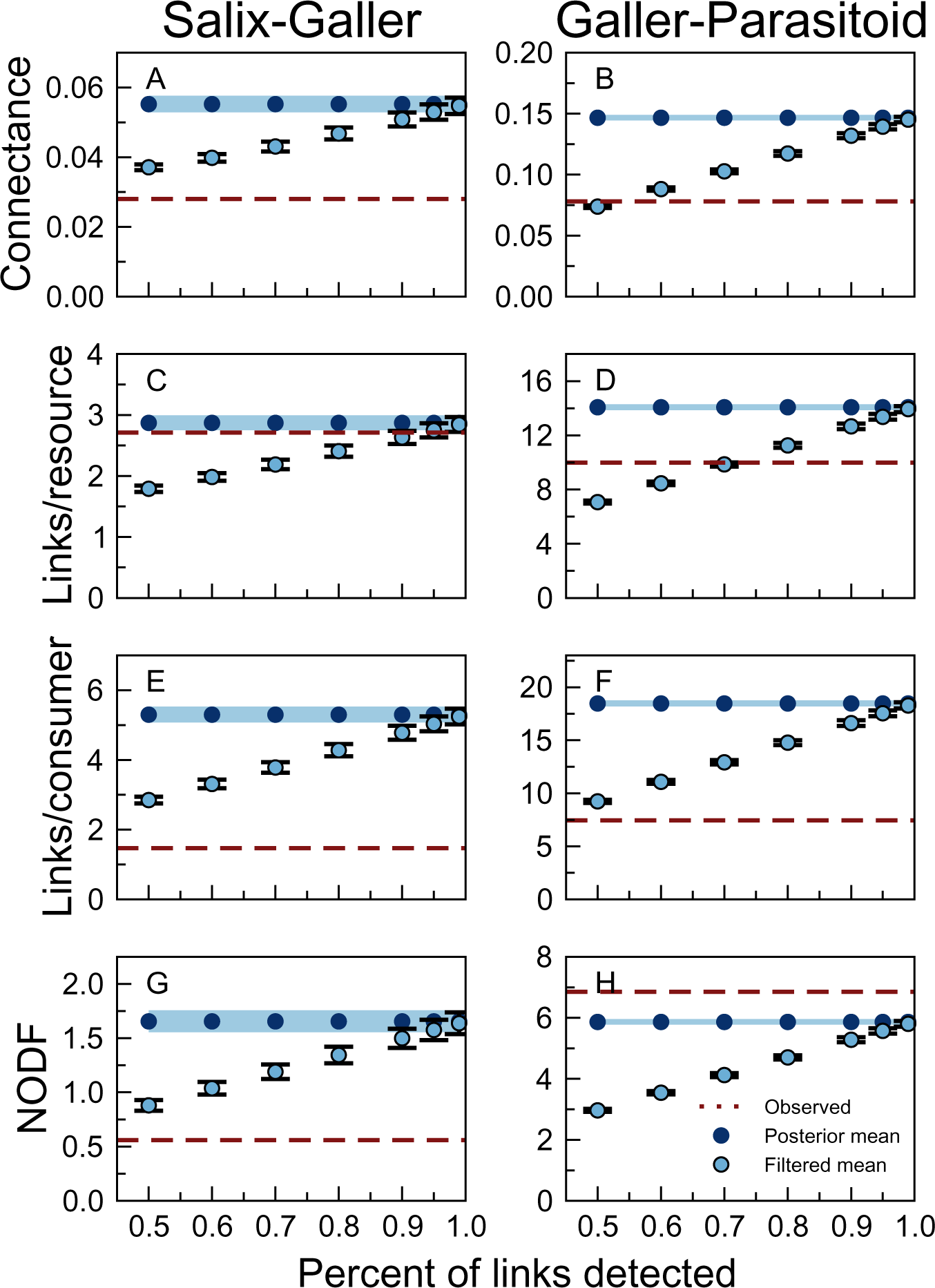
Here we show the mean connectance, links per resource (*Salix* in the *Salix*-galler networks and gallers in the galler-natural enemy networks), links per consumer, and nestedness (NODF) for networks assembled using posterior distributions based on a single sub-network in the Kopelke et al. (2017) dataset (Zillis). We created 100 “posterior-sampling” networks and then, for each of these, created 100 “detection-filter” networks by randomly sampling 50%-99% of the interactions included in the posterior-sampling network. This simulates imperfect detection of interactions in the field. Each point represents the mean network property (e.g., connectance) obtained from a set of 100 detection-filter networks, plotted against the value of the network property in the posterior-sampling network used to create the detection-filter networks. For each property and both network types, the posterior-sampling networks cover a relatively small range of network properties than the range covered by networks with varying detection probabilities. The value of each property decreases with the proportion of links included in the detection-filter networks.

Considering the galler-natural enemy networks, the connectance, mean links per galler, and mean links per natural enemy were also much lower in the observed web (C=0.078, *L_galler_*=9.99, and *L_naturalenemy_*=7.45, respectively) than in the posterior webs (0.186 ≤ C ≤ 0.198, 13.4 ≤ *L_galler_* ≤ 14.6, and 23.4 ≤ *L_naturalenemy_* ≤ 25.0). When the detection probability was relatively low (i.e., 50%), however, the properties of randomised networks became similar to those in the observed webs (Fig. 5B,D,F). Nestedness was higher in the observed network (NODF=6.85) than in the posterior webs (6.31 ≤ NODF ≤ 6.82; Fig. 5H); in this case, the stronger the detection filter the farther apart were the observed and posterior webs.

## Conclusions / recommendations

Real interaction networks vary over several dimensions (Kitching & Kitching, 1987; Olesen et al., 2011; Pires et al., 2011a; Baiser et al., 2012; Fodrie et al., 2015; Novak & Tinker, 2015) and to capture this variation we must turn from static descriptions of network structure to probabilistic descriptions. In this study, we have developed the analytical tools to capture the uncertainty in the estimation of pairwise interactions and a conceptual framework for its individual components: interaction uncertainty, process uncertainty, and detection uncertainty. Using this framework leads us to offer tangible recommendations for improved descriptors of ecological interactions. First, our analyses point to detection uncertainty as a major contributor to overall uncertainty of is establishing the absence of interaction. To counter this and establish true absences of
interactions requires comparatively large sample size on the order of 30-50 observations per species pair. Second, where such extensive sampling is not feasible, researchers should still acknowledge the varying levels of confidence surrounding the presence or absence of interactions between different pairs of species. Including the *n* and *k* values for each interaction will clearly indicate which unobserved interactions are most likely to be observed with further sampling and which estimates are more reliable. Third, the uncertainty around interactions (especially interactions that were not observed) should be incorporated in calculations of network properties like connectance or nestedness. Re-sampling networks based on a probabilistic understanding of networks is straightforward and gives distributions for network properties rather than point estimates. This not only acknowledges the fact that interactions vary over time and space but will also facilitate comparisons between networks. With confidence intervals around network metrics, we can not only say that one network is more connected than another but also whether the networks are more different than we would expect based on imperfect sampling of interactions. To facilitate these recommendations, we provide all code used in this paper in the supplementary material.

## Acknowledgements

The authors thank Daniel Cartensen for fruitful discussion of the ideas in this manuscript. We also thank Kévin Cazalles for providing feedback on the manuscript. The authors also appreciate support from the Swedish Research Council (VR) for grant #2016-06872 (to TR). Additional funding was provided by a Formas grant (#942-2015-1262) to AE.

